# PROX1 transcription factor is a master regulator of myogenic and oncogenic features of rhabdomyosarcoma

**DOI:** 10.1101/2020.04.19.045989

**Authors:** Nebeyu Yosef Gizaw, Pauliina Kallio, Tatjana Punger, Matias Kinnunen, Erika Gucciardo, Kaisa Lehti, Caj Haglund, Tom Böhling, Markku Varjosalo, Mika Sampo, Kari Alitalo, Riikka Kivelä

**Affiliations:** Stem Cells and Metabolism Research Program, Research Programs Unit, Faculty of Medicine, University of Helsinki, Helsinki, Finland; Translational Cancer Medicine Research Program and Center of Excellence, Research Programs Unit, Faculty of Medicine, University of Helsinki, Helsinki, Finland; Institute of Biotechnology and Helsinki Proteomics Unit, HiLIFE, University of Helsinki, Helsinki, Finland; Individualized Drug Therapy Research Program, Research Programs Unit, Faculty of Medicine, University of Helsinki, Helsinki, Finland; Department of Microbiology, Tumor and Cell Biology, Karolinska Institutet, Stockholm, Sweden; Department of Surgery, University of Helsinki and Helsinki University Hospital, Helsinki, Finland; Department of Pathology, University of Helsinki and Helsinki University Hospital, Helsinki, Finland; Department of Pathology, University of Helsinki and Department of Pathology / HUSLAB; Wihuri Research Institute, Helsinki, Finland

**Keywords:** cancer, PROX1, histone deacetylation, HDAC, metabolism

## Abstract

Rhabdomyosarcoma (RMS) is an aggressive pediatric soft tissue cancer in need for novel therapies. Here we show that the PROX1 transcription factor, which is essential for normal myoblast differentiation, is highly expressed in RMS tumors. We demonstrate that PROX1 is needed for RMS cell stemness and growth *in vitro*, and for RMS tumor formation in mouse xenograft models. In addition, we unveil that PROX1 is an essential for myogenic properties in RMS. PROX1 depletion reprogrammed the RMS transcriptome to resemble benign mesenchymal stem cells and repressed many of the previously identified RMS effector transcripts and myogenic genes. By using proximity labeling and mass spectrometry, we found that PROX1 interacts with the NuRD and CoREST complexes containing class I HDACs. Our studies reveal a major role of PROX1-HDAC interaction in RMS and give insights that inhibiting this interaction could be a promising therapeutic approach.

## Introduction

Rhabdomyosarcoma is the most common, highly aggressive pediatric soft tissue sarcoma, accounting for about 50 percent childhood and three percent of adult soft tissue sarcomas (Kashi et al., 2015). Two main types of RMS are distinguished histologically: embryonal (ERMS), which represents approximately 60% of all RMS and alveolar (AR, which accounts for about 25% of RMS (Newton et al., 1988). About 80% of ARMS are typically associated with pathogenic chromosomal translocation resulting in the expression of a PAX3-FOXO1 or PAX7-FOXO1 fusion protein (Davis, D’Cruz, Lovell, Biegel, & Barr, 1994). The remaining 20 % lacking this translocation are classified as fusion negative, and their clinical outcome is more similar to ERMS. Thus, in clinics RMS is now often classified being either fusion positive or fusion negative (FP-RMS and FN-RMS) (Williamson et al., 2010).

In about 25 % of patients, the disease is already metastasized during the diagnosis, and the 5-year survival in this case is only 20-30 % (Rudzinski el al., 2017). Another challenge is the recurrence of the disease as this is common among RMS patients. The treatment of RMS includes surgical removal of the tumor, radiation and cytostatics. As most of the RMS patients are children, unspecific treatments such as cytostatic treatment, produce long-term side effects including for example heart problems and an increased risk for another malignancies (Kashi et al., 2015). Despite intensive research, high-risk RMS treatment has not improved substantially for several decades, which emphasizes the need to further uncover the molecular mechanisms regulating the development and growth of RMS.

RMS tumors have been suggested to arise from the muscle stem cells (i.e. satellite cells), which fail to differentiate into mature skeletal muscle fibers (Tiffin et al., 2003). During myogenesis, the temporal expression of myogenic regulatory factors MYOD1, MYF5, MYF6, and myogenin drive differentiation and a terminal cell-cycle exit (Buckingham & Rigby, 2014). RMS cells express most of these factors, yet they fail to execute terminal differentiation (Tenente et al., 2017). It has also been suggested that RMS can arise from mesenchymal progenitor cells that reside in several non-muscular tissues (Charytonowicz et al., 2009). Recently, animal models for FP-RMS and FN-RMS demonstrated the origin of RMS from both myogenic and non-myogenic precursors, both acquiring myogenic features during tumor development (Yohe et al., 2019). Thus, identification of the cells that can give rise to RMS still remains one of the most challenging and thrilling question to be answered in RMS development (Kashi et al., 2015). This emphasizes the need to uncover the common molecular mechanisms regulating the myogenic phenotype in RMS independent of the origin.

Our previous work has shown that the prospero-related homeobox 1 (PROX1) factor is expressed in muscle stem cells, and that it is essential for myoblast differentiation and slow muscle fiber type characteristics (Kivela et al., 2016). PROX1 is highly conserved among vertebrates and it is essential for lymphatic endothelial cell differentiation and for the development of eye, liver and heart (Elsir et al., 2012). Both tumor suppressive and oncogenic properties have been attributed to PROX1, as its altered expression levels have been found in a variety of human cancers, such as colorectal cancer, Kaposi’s sarcoma, pancreatic and hepatocellular carcinoma (Elsir et al., 2012). This indicates that PROX1 can function either as an oncogene or a tumor suppressor, depending on the tissue and cancer type.

As PROX1 has been shown to maintain cancer stem cells in colorectal cancer by recruiting the NuRD complex (Hogstrom et al., 2018), and as our previous findings have demonstrated PROX1 to be an essential factor regulating skeletal muscle phenotype and myoblast differentiation, we hypothesized that PROX1 could also play a role in the development and progression in RMS. To study this, we have used biobanked human tumor samples, alveolar and embryonal RMS cell lines, primary patient cells, gene silencing, transcriptomics, proteomics analysis, colony and rhabdosphere formation assays and tumor xenograft mouse models. Based on the findings, we show that PROX1 is essential for the myogenic phenotype, stemness and growth of RMS.

## Results

### PROX1 is highly expressed in rhabdomyosarcoma

To gain insight into PROX1 expression in RMS, we analysed RNA expression profiles in various patient cohorts and sarcoma subtypes using publicly available datasets deposited in Gene Expression Omnibus (https://www.ncbi.nlm.nih.gov/geo/), Oncomine (https://www.oncomine.org/resource/login.html) and cBioportal (https://www.cbioportal.org/). Analysis of RNA sequencing data from primary patient samples (GEO: GSE108022) (Shern et al., 2014) showed that PROX1 expression in RMS is higher than in healthy skeletal muscle in 50% of the FN-RMS cases (n= 33/66) and in almost all of the FP-RMS subtypes (n=34/35) (**Figure 1A**). Based on microarray data (GEO: GSE22520) Prox1 expression was also higher in both ERMS and ARMS mouse models than in healthy skeletal muscle (Abraham et al., 2014; Rubin et al., 2011) (**Figure 1B**). The comparison between different sarcoma types (GEO: GSE2553) indicated that high PROX1 expression is relatively specific to RMS, as there were only few Ewing sarcomas and synovial cell sarcomas demonstrating high PROX1 expression (Baird et al., 2005) (**Figure 1C**). PROX1 RNA and protein was also highly expressed in all studied ERMS (RD, RH36, and RMS) and ARMS (KLHEL1 and RH30) cell lines, when compared to healthy human myoblasts (**Figure 1D and 1E**). Immunohistochemistry of 155 RMS tumor samples from the Helsinki Biobank revealed high PROX1 protein expression in the majority of primary and metastatic RMS tumor samples (n = 142/155). In particular, intense PROX1 staining was found in almost all of the primary tumor samples either within the entire tumor nuclei or in dispersed pattern (n = 24/24 ERMS and n = 20/21 ARMS), (**Figure 1F and 1G**). Similarly, immunofluorescent PROX1 staining was observed in the nuclei of RD cells, representing one of the most commonly used cell lines in RMS research (Hinson et al., 2013) (**Figure 1F**).

**Figure 1.**
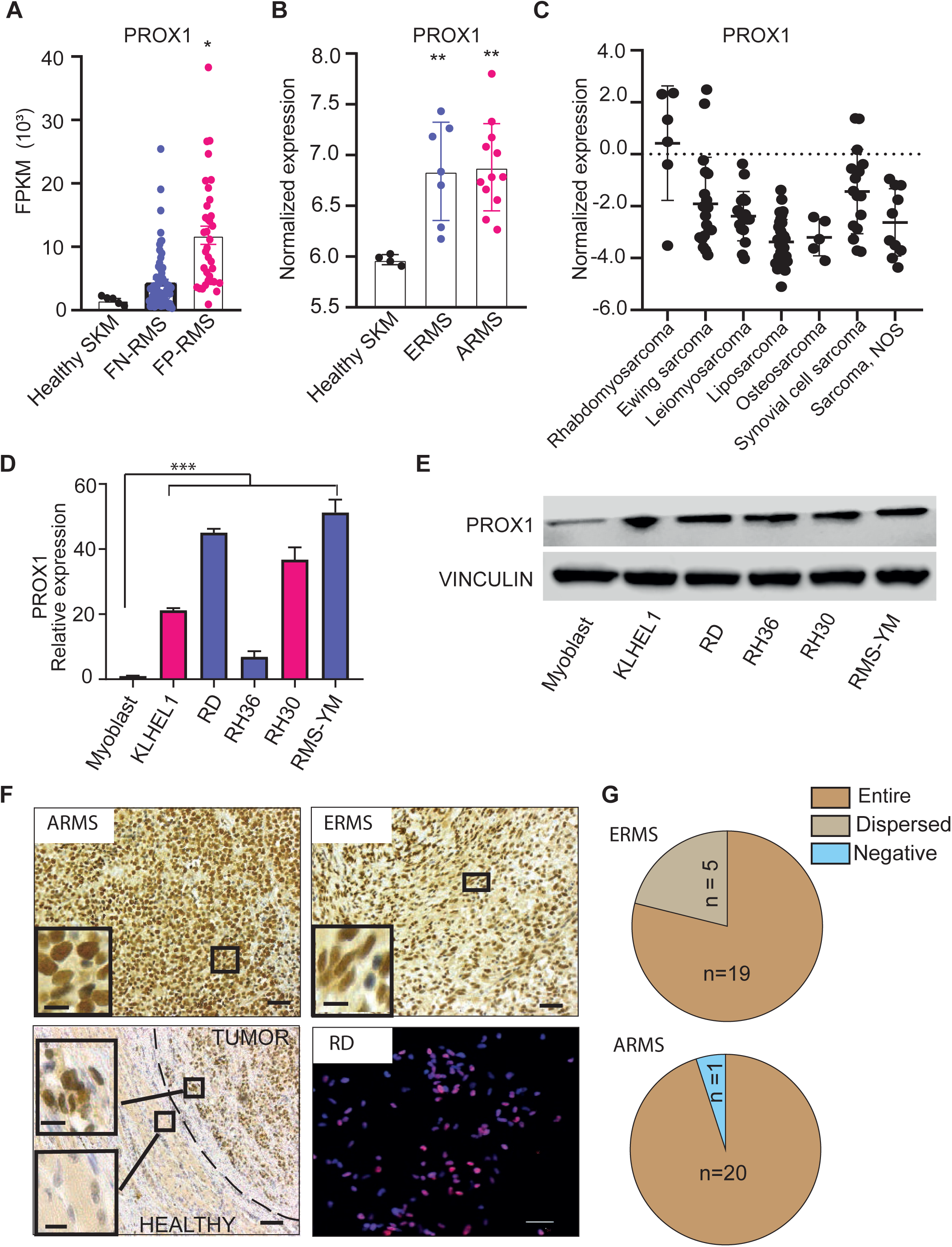
High PROX1 expression in primary RMS tumors and metastases. (**A**) Normalized mRNA expression (FPKM) for PROX1 in normal muscle (Healthy SKM), fusion negative and fusion positive RMS patient sample (GSE108022, Adj *p < 0.05). (**B**) mRNA expression of PROX1 in healthy mouse skeletal muscle and in embryonal (ERMS) and alveolar (ARMS) experimental tumors (GSE22520 microarray data, Adj **p < 0.01). (**C**) Microarray data depicting PROX1 expression in different sarcomas (GSE2553). (**D**) Real-time qPCR analysis of PROX1 in RMS fusion negative (RD, RH36 and RMS-YM) and fusion positive (KLHEL1 and RH 30) cell lines compared to healthy human myoblasts (***p < 0.001). (**E**) Western blot analysis of PROX1 in RMS cell lines and in healthy human myoblasts. Vinculin was used as a loading control. (**F**) Immunohistochemical staining of PROX1 in human tumor samples from Helsinki Biobank. Both ARMS and ERMS patient samples show strong nuclear PROX1 expression. The lower left panel shows the difference in PROX1 expression between the tumor and the healthy muscle. Dashed line marks the tumor boundary. In lower right panel RD cells are stained for PROX1 (red) and nuclei (blue). Scale bars for full image 50µM and for the magnified inset 10 µM (**G**) Classification of PROX1 expression pattern in the primary RMS tumor samples from the Helsinki Biobank. The data used in the panels a-c was retrieved from the NIH Gene Expression Omnibus. Data is presented as mean±SEM

### PROX1 is required for RMS cell growth

To study the functional significance of PROX1 in RMS, we silenced PROX1 RNA expression by using two independent lentiviral short hairpin RNAs (shRNAs) both in the ERMS cell line RD and in the KLHEL1 cell line that we derived from an ARMS patient. Four days after the transduction, the PROX1 silenced RD and KLHEL1 cells had developed a rounded morphology that was distinct from the spindle shaped form observed in the control vector (shSCR) transduced cells (**Figure 2A and 2B**). Analysis of 5-ethynyl-2’-deoxyuridine (EdU) incorporation showed significantly reduced proliferation in the PROX1 silenced RD and KLHEL1 cells (**Figure 2C and 2D**). The colony formation assay (Li et al., 2013) used to evaluate the stemness potential of the RMS cells revealed that the PROX1 silenced RD cells were unable to form any colonies and very few were observed in cultures of the PROX1 silenced KLHEL1 cells (**Figure 2E and 2F**). Further, in the 3D spheroid assay, which is a common tool to measure tumor self-renewal capacity *in vitro* (Pastrana et al., 2011), the PROX1 silenced RD and KLHEL1 cells formed significantly less rhabdospheres than the shSCR transduced cells (**Figure 2G – 2L**). Importantly, the spheroids formed from the shPROX1-GFP transfected cells were mostly GFP negative, indicating that they were derived from escape clones, which had lost the PROX1 silencing construct, whereas the spheroids formed from control shSCR-GFP treated cells retained GFP positivity (**Figure 2I and 2J.)** These data demonstrate that PROX is essential for the stemness properties of RMS cells regardless of the RMS subtype.

**Figure 2.**
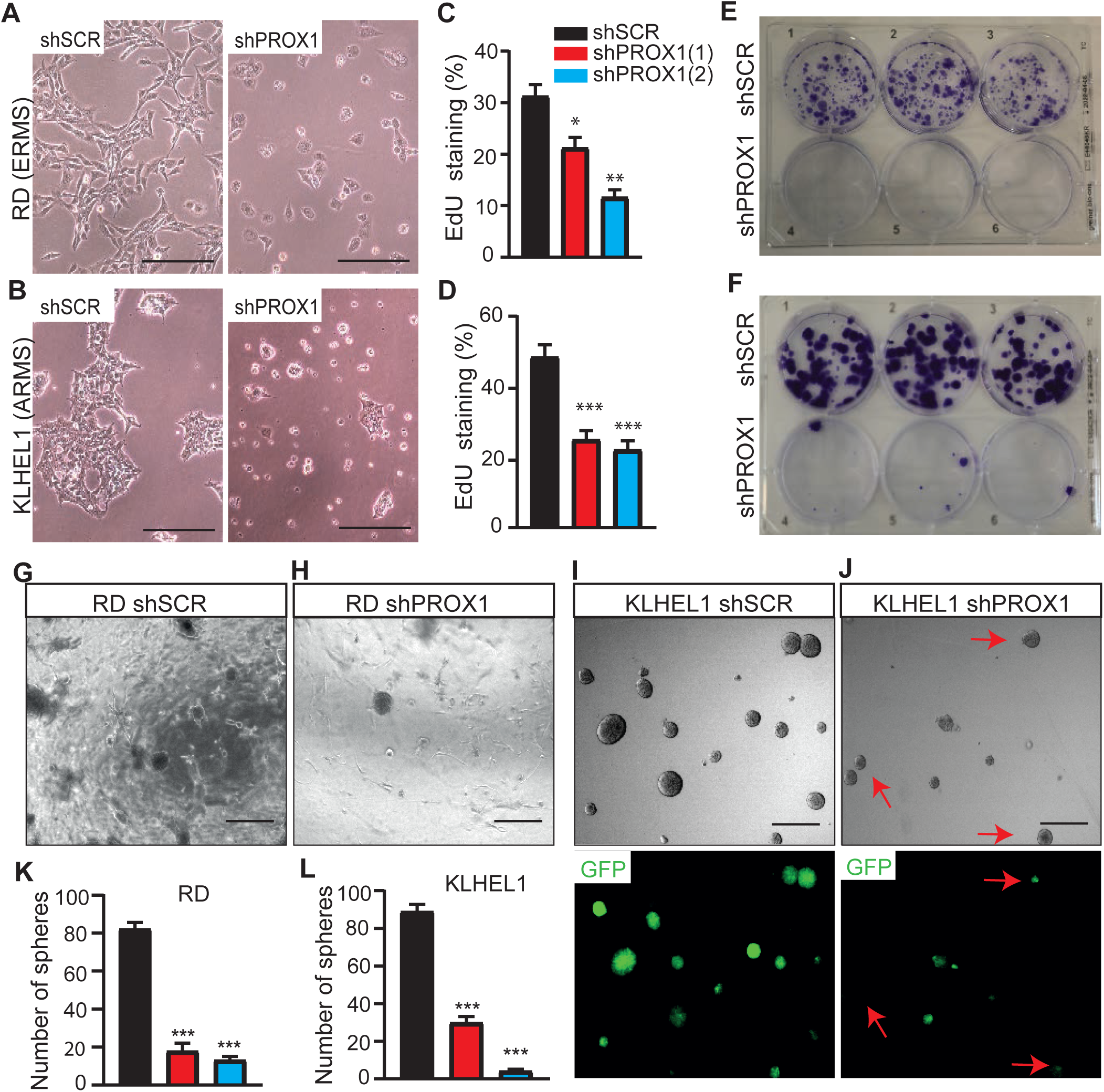
PROX1 promote RMS cell proliferation and is required for stemness. (**A-B**) Bright field images depicting morphological changes in RMS cells infected with shSCR and shPROX1 at 4 days post infection (**A**) for RD cells (**B**) for KLHEL1. Scale bars 50µM. (**C-D**) EdU incorporation analysis for proliferation rate in shSCR and shPROX1 RD (**C**) and KLHEL1 (D) cells. (**E-F**) Colony formation assay in shSCR or shPROX1 treated RD (**E**) and KLHEL1 (F) cells. Cells were fixed and stained with crystal violet 9 days after seeding. (**G – J**) 3D rhabdosphere formation in shSCR and shPROX1 RD and KLHEL cells. Scale bars 500µM. (**I-J**) Representative images showing that rhabdospheres formed from shPROX1 KLHEL1 cells consisted mainly from escape clones that were devoid of shPROX1-GFP infection. (**K-L**) Quantification of rhabdospheres formation (2000 cells for RD and 500 cells for KLHEL1 were plated per well (n = 4 for each cell line) and rhabdospheres were quantified at day nine). Two independent shPROX1 constructs were used. *P < 0.05, **P <0.01 and ***P<0.001. Data is presented as mean±SEM.

### PROX1 is essential for the growth of RMS tumor xenografts

After obtaining strong evidence that PROX1 regulates RMS cell growth and rhabdosphere formation *in vitro*, we next assessed whether PROX1 has tumor propagating potential also *in vivo*. RD cells were transduced with lentiviral vector encoding shSCR and GFP or shPROX1 and GFP, which resulted in approximately 50-60% reduction in PROX1 RNA expression (**Figure 3A**). The shPROX1 and shSCR control cells were transplanted subcutaneously into the left and right flanks of NOD/SCID/IL2rg null (NSG) mice, respectively. Serial tumor volume measurements showed that the PROX1 silenced tumors had markedly delayed tumor growth and reduced tumor volume in comparison with the shSCR tumors (**Figure 3B**). The weights of PROX1 silenced tumors were significantly lower than their controls at the end of the experiment (**Figure 3C-E**). qPCR analysis of the excised tumors showed that, despite its silencing, PROX1 RNA was re-expressed to a similar degree in both shPROX1 and shSCR tumors. Furthermore, unlike in the shSCR tumors, only few GFP-positive cells were found in shPROX1 tumors (**Figure 3F-G**). This indicated that escape clones were responsible for the growth of the shPROX1 tumors, similarly as observed in the rhabdosphere assays *in vitro*. Similar results were obtained with another PROX1 silencing construct and when using a shorter follow-up time (**Supplement Figure1**). Together, these results demonstrate that PROX1 is an essential regulator of RMS tumor growth both *in vitro and in vivo*.

**Figure 3.**
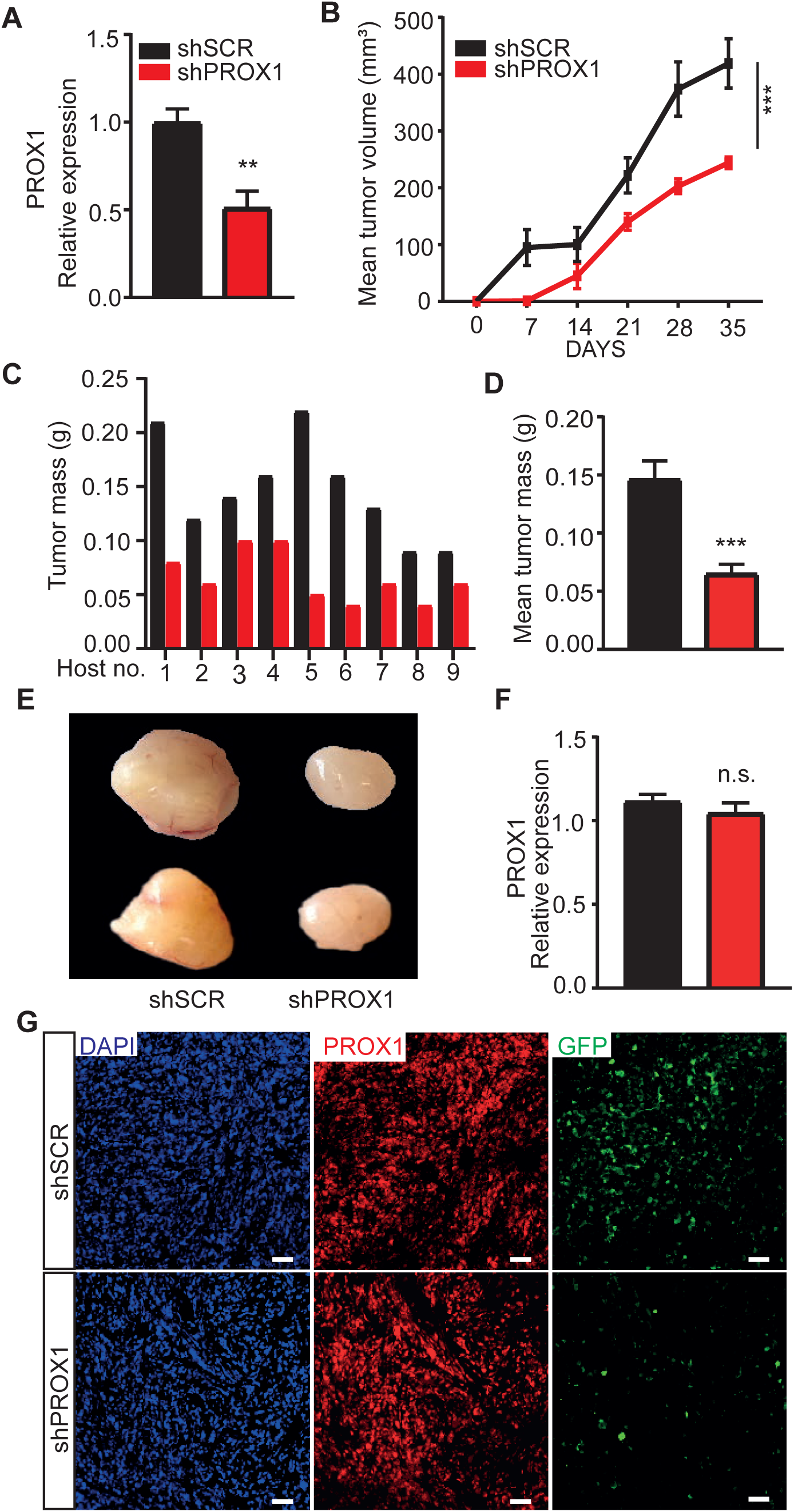
PROX1 is required for tumor xenograft growth. (**A**) RT-qPCR analysis of the PROX1 relative expression shPROX1 RD cells before tumor implantation. (**B**) Quantification of tumor growth based on tumor volume (mm^3^) for shSCR and shPROX1 RD cells injected into NSG mice left and right flank region, respectively (n = 9 per group). (**C and D**) Comparison of individual tumor masses from shSCR or shPROX1 tumors in same host (C) and average tumor mass (**D**) at the end of the experiment (day 35). (**E**) Representative images of shSCR (left) and shPROX1 (right) xenograft tumors. (**F**) RT-qPCR analysis of PROX1 expression in xenograft tumors from shSCR or shPROX1 RD cells (**G**). Tumor sections from shSCR-GFP and shPROX1-GFP stained for PROX1 (red), GFP (green) and nuclei (DAPI, blue). Note that the shPROX1 tumors had regained PROX1 expression and only few cells were GFP positive indicating that the tumors were formed from escape clones. **P <0.01 and ***P<0.001. Data is presented as mean±SEM.

### PROX is a master regulator of myogenic and malignant sarcoma transcriptome in RMS cells

To decode the molecular basis of PROX1 dependent regulation of RMS tumor growth, we performed RNA sequencing (RNA-seq) analysis of four biological replicates of PROX1 silenced (75% deletion) and control RD cells. Principal component analysis (PCA) and hierarchical clustering of the RNA-seq data showed that the control and the PROX1 silenced cells fell into two distinct groups (**Figure 4A and 4B**). Analysis of differentially expressed genes (DEG), using a statistical significance cut-off at FDR < 0.05 and biological significance cutoff at ≥ 1.5-fold, showed 461 upregulated and 433 downregulated genes in the shPROX1 RD cells **(Table S1)** and 1141 up- and 1119 downregulated genes when using FDR < 0.05 and log2 fold-change ≤ −0.25 and ≥ 0.25 (**Figure 4C).** Gene ontology analysis revealed that cell-cell adhesion, angiogenesis, and cell-matrix adhesion were the most significantly enriched GO terms among the upregulated DEGs (**Figure 4D**), and skeletal muscle contraction and muscle filament gliding were enriched among the downregulated DEGs (**Figure 4E**). Intriguingly, these GO terms have been previously shown to be among the top biological processes that shift, in an opposite manner, when non-myogenic mesenchymal cells were driven into FP-RMS (Ren et al., 2008), and in FN-RMS (Drummond et al., 2018). Importantly, GSEA analysis revealed that the gene sets altered in mesenchymal stem cells after forced expression of PAX3-FKHR(FOXO1) fusion gene to drive RMS tumorigenesis (Ren et al., 2008), were changed in an opposite direction by PROX1 silencing, i.e. towards a benign mesenchymal cell phenotype. This demonstrates that PROX1 and the PAX3-FOXO1 fusion gene that drives RMS regulate an overlapping set of transcripts associated with malignancy and myogenic transcriptome (**Figure 4F and 4G**). Several genes regulating the mesenchymal stem cell (MSC) phenotype were found to be significantly induced by PROX1 silencing (e.g. THY1, NT5E, CD44) and genes related to myogenic features were repressed (e.g. MYOG, MEF2C, MYH7, MYH8, MYH3, MYL1, CAV3, TTN, TNNT1, TNNC1, TNNC2, PAX7, ACTC1) (**Supplement Table 1**). GSEA also revealed that hallmark genes for myogenesis were highly downregulated in PROX1 silenced RD cells (**Figure 4H**). To consolidate further, whether PROX1 silenced RD cells gain mesenchymal stem cells characteristics, we stained for the MSC marker NT5E (CD73) (Dominici et al., 2006). In line with transcriptomic data, the expression of CD73 was highly increased in PROX1 silenced RD cells (**Figure 4I and 4J**). The GSEA analysis also showed that PROX1 silencing induced the expression of transcripts, which were downregulated in mesenchymal stem cells and in the UET-13 mesenchymal progenitor cells after ectopic expression of EWS/FLI or EWS/ERG fusion genes that drive Ewing’s family tumors (EFT) (Miyagawa et al., 2008) (Riggi et al., 2008) (**Supplement Figure 2A and 2B**). This suggests that PROX1 silencing can, at least partially, revert the malignant transcriptomic phenotype of RMS and EFT.

**Figure 4.**
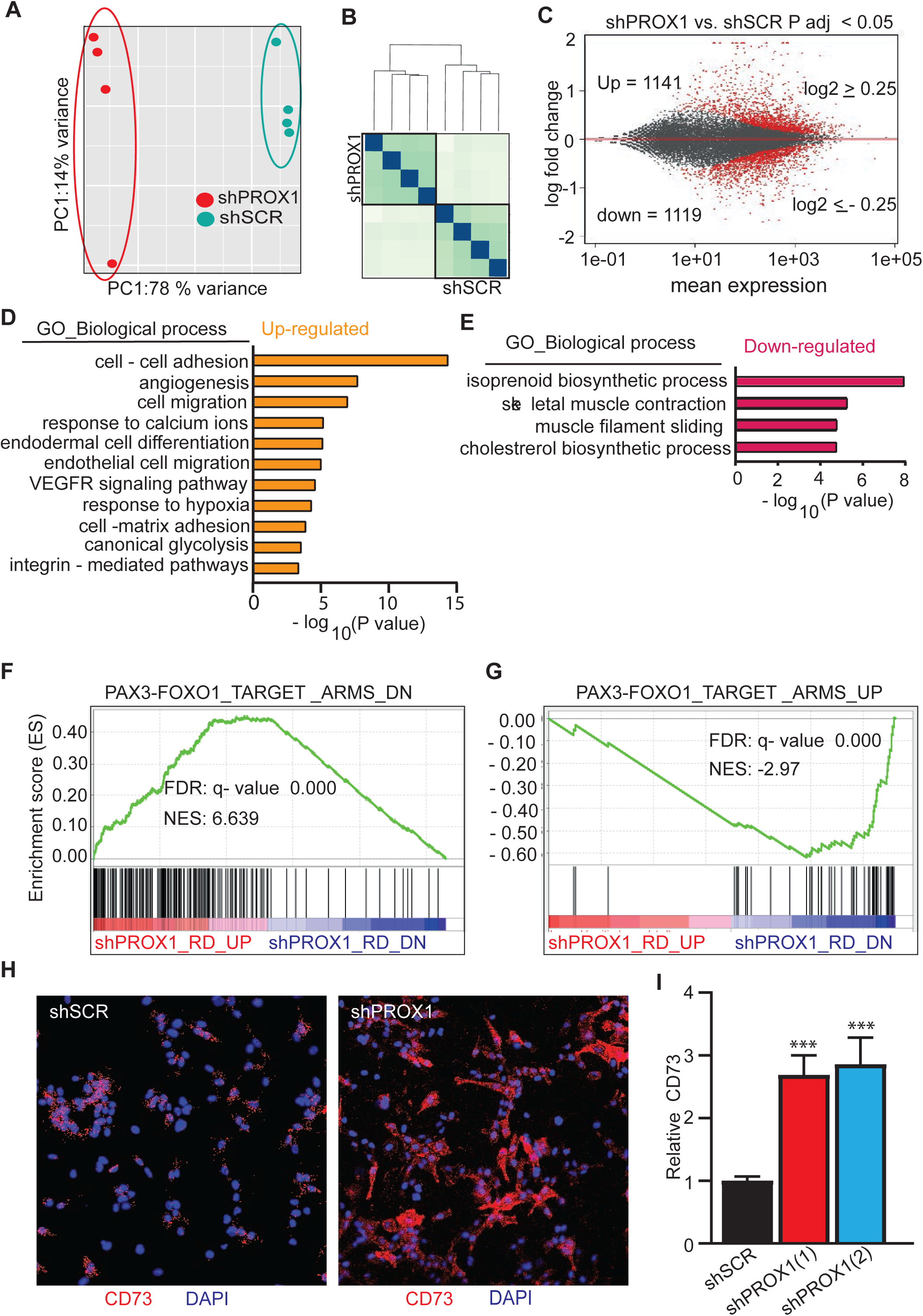
PROX1 is a potent regulator of malignant transcriptome in sarcoma cells. Global gene expression analysis of PROX1 silenced RD cells by RNA sequencing (**A-I**). (**A**) Principal components analysis of the variance in between samples and (**B**) sample distance analysis of RNA-seq samples (N=4+4). (**C**) MA plot representing the differentially expressed genes in red (P adj < 0.05, log2 FC cut-off 0.25) in PROX1 silenced compared to control RD cells, including 1141 upregulated and 1119 downregulated genes. (**D and E**) Gene Ontology analysis showing the most significant functional categories among the upregulated genes (**D**) and the downregulated genes (**E**) in PROX1 silenced cells. (**F and G**) Gene Set Enrichment Analysis plots for gene sets overlapping with PROX1 driven differentially expressed genes in RD cells. Enrichment of downregulated genes (**F**) and upregulated genes (**G**) in mesenchymal stem cells overexpressing the PAX3-FOXO1 fusion gene. NES, normalized enrichment score; FDR, false discovery rate. Each black bar represents a single gene within a gene set. (**H**) Immunofluorescence staining for CD73 (red) and nuclei (blue) in scramble shRNA (shSCR) and PROX1 shRNA (shPROX1) RD cells. (**I**) Quantification of CD73 fluorescence intensity in shRNA - treated RD cells (***p < 0.001). Error bars represent SE. Data is presented as mean±SEM

### PROX1 regulates the expression of the major pro-oncogenic genes identified in RMS

From the RNAseq data, we analyzed the expression of the oncogenes that have previously been shown to be upregulated, and characterized as tumorigenic, in RMS. Several such genes were found to be significantly repressed by PROX1 silencing: FGFR4, MYL4 (Khan et al., 2001), IGFBP5, IGF2 (El-Badry et al., 1990; Khan et al., 1999), MYOG (Gryder et al., 2017), MYCN (Williamson et al., 2005), RASSF4 (Crose et al., 2014), VANGL2 (Hayes et al., 2018), Hey1 (Belyea et al., 2011) and NOTCH3 (De Salvo et al., 2014) (**Figure 5A**). Correlation analysis of these genes with PROX1 using RMS tumor RNA expression data in MediSapiens (n = 49 tumors, http://ist.medisapiens.com/) showed significant correlation between PROX1 and six of these genes (**Supplement Figure 3**), of which one of the strongest correlations was found between PROX1 and FGFR4 (**Figure 5B**). A population of high PROX1 and FGFR4 co-expressing colorectal cancer cells were previously shown to form aggressive tumors and to be resistant to cell arrest (Petrova et al., 2008). The correlation was validated by qPCR in PROX1 silenced RD and KLHEL1 cells (**Figure 5C**). To examine if FGFR4 silencing would phenocopy the effects of PROX1 silencing, we used two independent shFGFR4 constructs (**Figure 5D**). The colony and rhabdosphere formation capacity of RD cells was significantly reduced in the FGFR4 silenced cells (**Figure 5E-H**). However, the overall effect was not as striking as in the PROX1 silenced cultures, which indicates that FGFR4 mediates the effects of PROX1 only partially.

**Figure 5.**
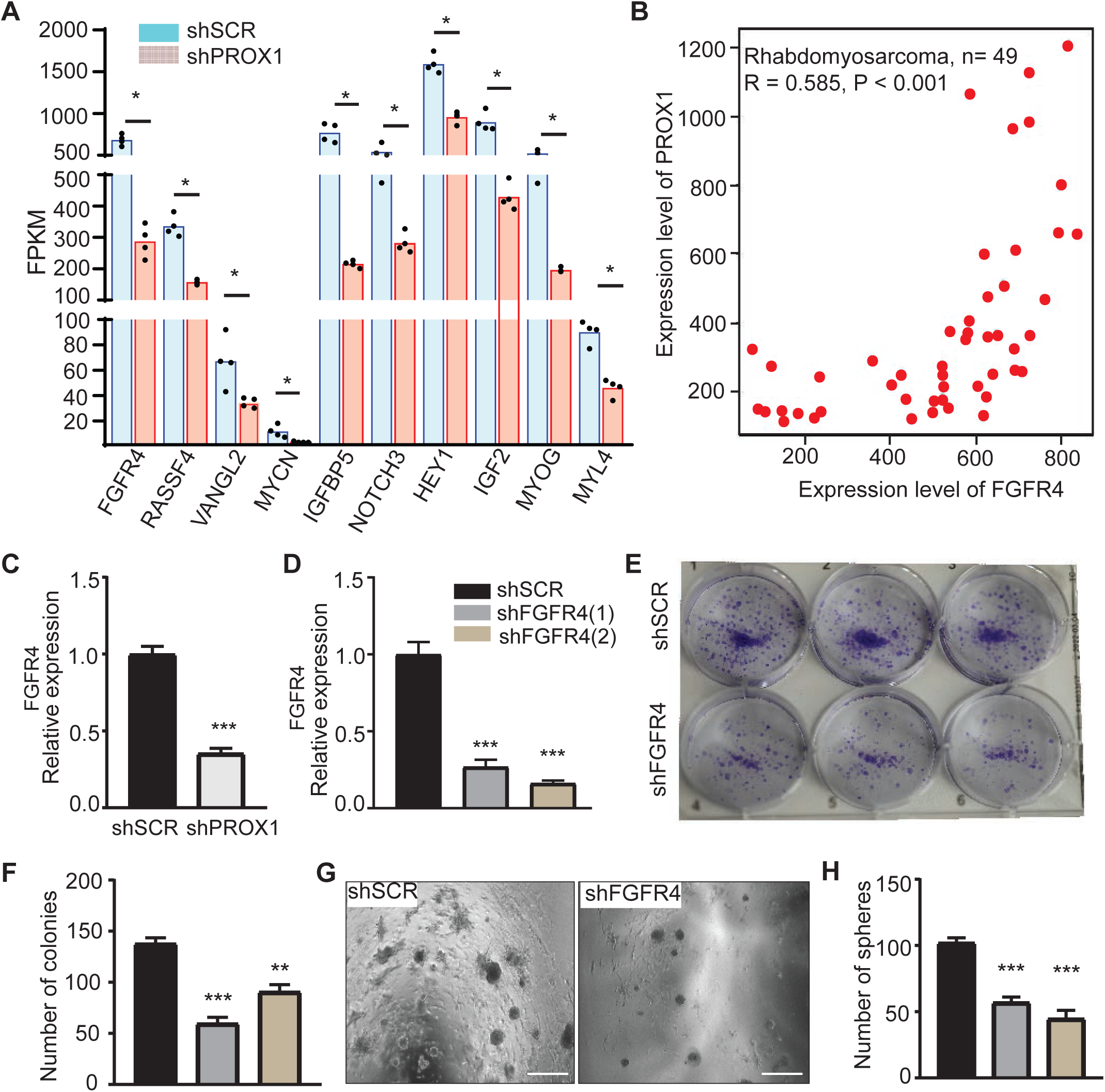
PROX1 is an upstream regulator of RMS associated pro-oncogenes. (**A**) Normalized expression values (FPKM) for RMS associated oncogenes in PROX1 silenced RD cells. (*Adj p < 0.05). (**B**) Correlation analysis of PROX1 and FGFR4 expression in RMS tumor RNA expression data. (**C**) RT-qPCR analysis of FGFR4 mRNA expression in PROX1 silenced vs control RD cells. (D –H) FGFR4 partially recapitulate the effect of PROX1 in colony and rhabdosphere formation in RD cells. (**D**) mRNA expression of FGFR4 in shSCR and in two independent shFGFR4 -treated RD cells. (**E**) Colony formation assay in shSCR and shFGFR4 -treated RD cells nine days after seeding and (**F**) quantification of colonies. (**G**) 3D rhabdosphere formation in shSCR and shFGFR4 RD cells nine days after seeding and (**H**) quantification of rhabdospheres. Scale bar 500 µM. ***P<0.001. Data is presented as mean±SEM.

### PROX1 regulates RMS cell metabolism

Since PROX1 has been shown to enhance metabolic adaptation in colon cancer cells (Ragusa et al., 2014), we performed KEGG pathway analysis of the RNAseq data, which revealed that citrate cycle, fatty acid (FA) metabolism and steroid synthesis were overrepresented among the DEGs downregulated in the PROX1 silenced cells (**Figure 6A and 6B**). To study the effects of PROX1 on RMS cell metabolism, we performed Seahorse bioenergetic analyses. We found that cellular oxygen consumption rate (OCR) was significantly decreased (**Figure 6C**) and extracellular acidification rate (ECAR) increased (**Figure 6D**) in the PROX1 silenced RD cells, indicating that PROX1 silencing shifted RD cell metabolism from oxidative to glycolytic. This is in line with the transcriptomic data showing increased expression of glycolysis genes and downregulation of genes related to fatty acid oxidation in PROX1 silenced RD cells. Glucose uptake did not differ between the PROX1 silenced and the control cells, but the uptake of free fatty acids was reduced by Prox1 silencing (**Figure 6E and 6F)**. However, despite the reduced FA uptake, the triglyceride content was significantly higher in the PROX1 silenced cells (**Figure 6G**), reflecting decreased FA oxidation and synthesis of structural and functional lipids.

**Figure 6.**
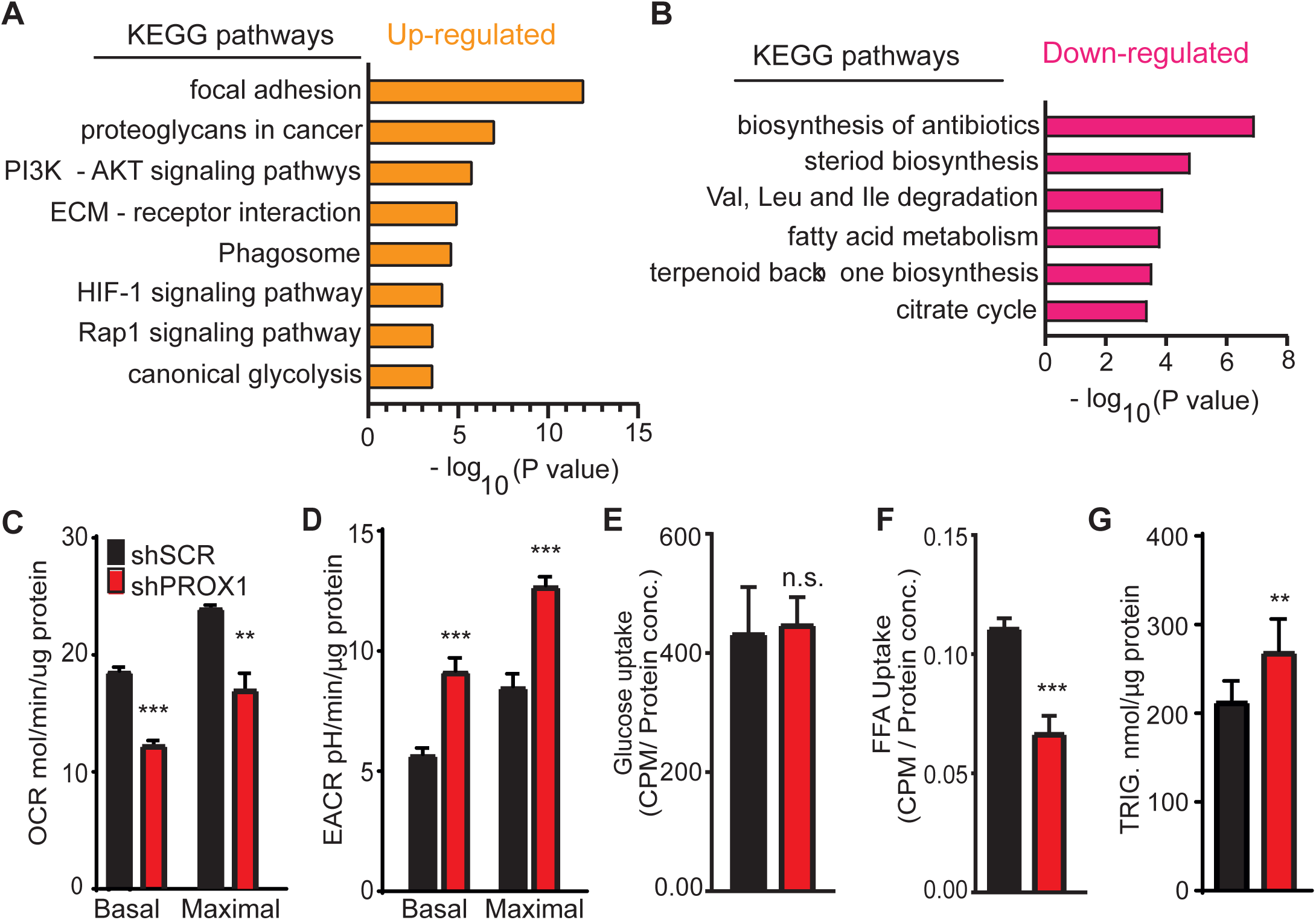
PROX1 regulates RMS cell metabolism. (**A and B**) Kyoto Encyclopedia of Genes and Genomes (KEGG) pathways enriched in the PROX1 silenced RD cells. Overrepresented pathways within the upregulated genes (**A**) and within the downregulated genes (**B**) plotted against the negative log of P value. (**C and D**) Bioenergetic analysis of shSCR and shPROX1 RD cells. (**C**) Basal and maximal oxygen consumption rate and (**D**) basal and maximal extracellular acidification rate (ECAR). (**E**) Glucose uptake and (**F**) fatty acid uptake in shSCR and shPROX1 RD cells. (G) Triglyceride content in shSCR and shPROX1 RD cells. **P <0.01 and ***P<0.001. Error bars represent SE. Data is presented as mean±SEM.

### PROX1 associates with CoREST and NuRD complexes to regulate its target genes

To investigate the mechanism of how PROX1 regulates its target genes, we performed BioID-mediated proximity labeling in RD cells expressing PROX1–BioID fusion protein. Mass spectrometry analysis of the biotinylated proteins revealed that PROX1 interacts with several proteins involved in chromatin remodeling. In particular, all components of the CoREST-deacetylase complex and nucleosome remodeling and deacetylase (NuRD) complex were identified as PROX1 binding partners (**Figure 7A and 7B, Table S2**). Interestingly, the class I histone deacetylases HDAC1, HDAC2 and HDAC3, but not other HDACs, were found to bind to PROX1. HDAC 1 and 2 are core components of the CoREST and NuRD complexes, whereas HDAC3 has been shown to interact with NCoR1 and NCoR2 in RMS cells (Phelps et al., 2016). Our data also show NCoR 1 and 2 as PROX1 binding partners in RD cells (**Supplement Table 2)**. Using co-immunoprecipitation, we validated that MTA1 and HDAC1, components of the NuRD and CoREST complexes, interact with PROX1 (**Figure 7C**).

**Figure 7.**
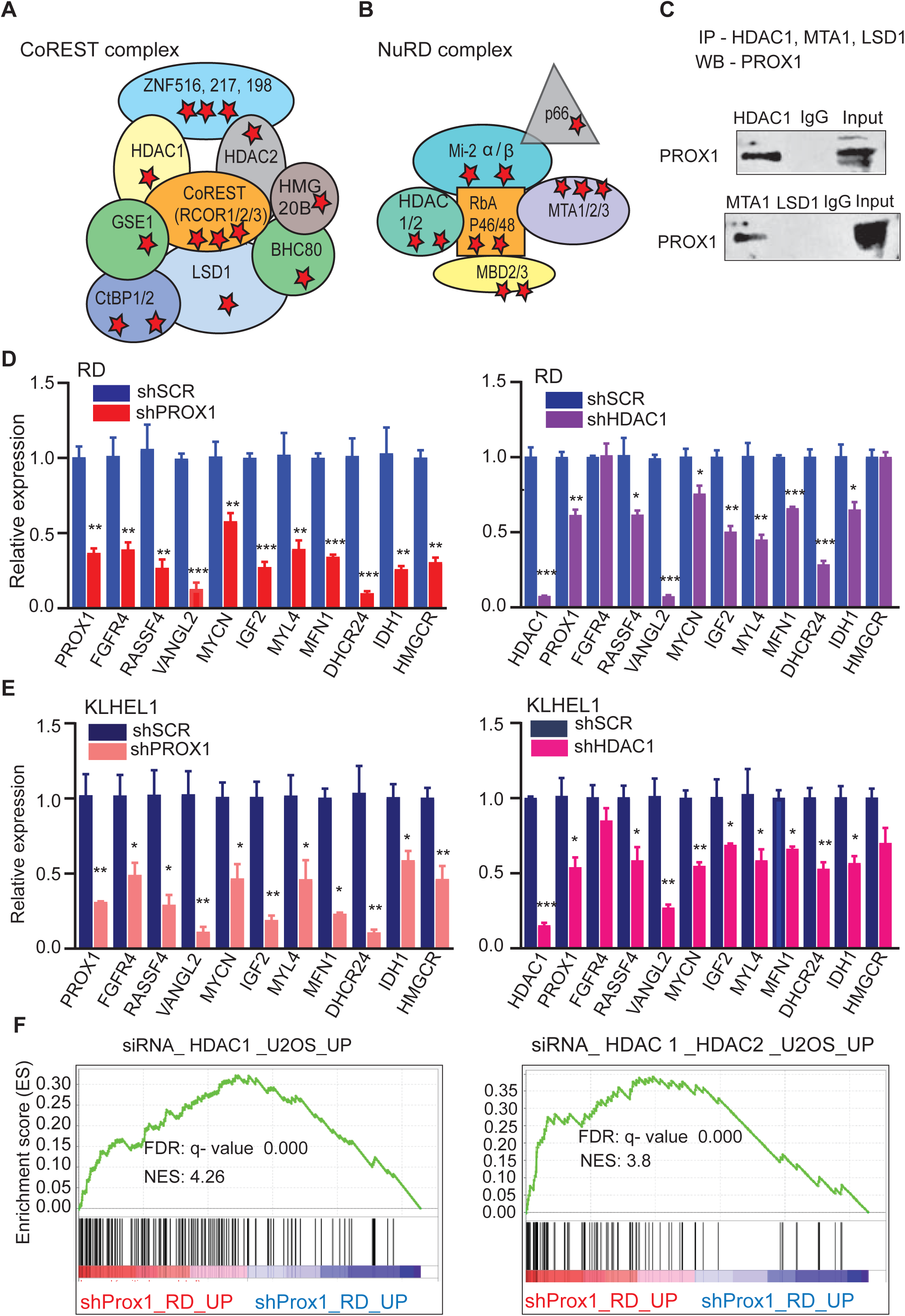
PROX1 co-operates with CoREST and NuRD complexes containing HDACs in RMS. (**A and B**) Schematic presentation of CoREST and NuRD which were identified as PROX1 binding partners in RD cells. Components of the complexes identified by PROX1 mediated proximity labeling and mass spectrometry in RD cells are marked with asterisk. Components of the CoREST (**A**) and NuRD (**B**) complexes. The red stars represent the identified PROX1-interacting components. (**C**) Co-immunoprecipitation and western blotting of PROX1 with HDAC1, LSD1 and MTA1 in RD cells. (**D and E**) qPCR analysis of RMS associated pro-oncogenes and metabolic genes in PROX1 (left) and HDAC1 (right) silenced RD cells (**D**) and KLHEL1 cells (**E**). (**F**) Gene Set Enrichment Analysis plots showing genes upregulated in HDAC1 (left), and in HDAC1 and HDAC2 (right) silenced U2OS osteosarcoma cells. Overlapping gene sets were upregulated by PROX1 silencing in RD cells. NES, normalized enrichment score; FDR, false discovery rate. NES, normalized enrichment score; FDR, false discovery rate. Each black bar represents a single gene within a gene set.

Next, we compared the effects of HDAC1 silencing to PROX1 silencing in RD and KLHEL1 cells. Similarly to PROX1 silencing, HDAC1 silencing repressed the expression of most of the RMS associated oncogenes, as well as PROX1 itself, in both RD and KLHEL1 cells (**Figure 7D and E**). Recent studies have demonstrated the therapeutic potential of HDAC inhibitors in cell cultures and in a pre-clinical RMS model (Bharathy et al., 2019; Bharathy et al., 2018). We studied the effects of vorinostat, a pan-HDAC inhibitor, using viability assays and demonstrated that its effect was enhanced by silencing of PROX1 (**Figure S4A**). Vorinostat also repressed mRNA and protein expression of PROX1 (**Figure S4B-D**), and attenuated colony formation in both RD and KLHEL1 cells (**Supplement Figure 4E**). Interestingly, the GSEA analysis identified an overlapping set of upregulated transcripts the HDAC1 or HDAC 1 and 2 silenced U2OS osteosarcoma cells (Senese et al., 2007) with PROX1 silenced RD cells (**Figure 7F**), indicating that PROX1 and HDAC1 regulate similar set of genes in sarcoma cells.

## Discussion

Our present work uncovers PROX1 transcription factor as a novel regulator of RMS phenotype and growth. PROX1 was strongly expressed in almost all FN- and FP-RMS primary tumors and their metastases in a large biobank cohort. PROX1 silencing in RMS cells repressed the hallmarks of myogenesis and decreased cell proliferation, clonogenic growth and rhabdosphere formation. Similar effect was observed in *in vivo* studies, where PROX1 silenced xenografts failed to develop tumors. Importantly, in addition to the loss of myogenic features, the expression of several genes previously associated with RMS development and growth was repressed in PROX1 deficient RMS cells, while the expression of mesenchymal stem cell genes was highly increased. Mechanistically, PROX1 interacts with CoREST and NuRD complexes containing class I HDACs.

During myoblast differentiation, PROX1 expression increases and it regulates the expression of other myogenic factors and components of the muscle contractile machinery (Kivela et al., 2016). In experimental RMS models, myogenic regulatory factors are expressed in the tumors regardless of their cells of origin (Tenente et al., 2017). Through chromatin acetylation, PAX3 – FOXO1 fusion gene has been shown to generate active super enhancers to drive MYOD1 and MYCN, which can then drive the expression of MYOG (Gryder et al., 2017). However, PAX-FOXO1 can activate myogenic determination and MYOG expression independent of MYOD (Davicioni et al., 2006; Zhang & Wang, 2007). A recent study demonstrated that activation of the hedgehog pathway by a constitutively activated form of Smo (SmoM2) in non-myogenic endothelial progenitor cells can lead to FN-RMS development (Drummond et al., 2018). Interestingly, we detected from their data that PROX1 was among the most upregulated genes during the tumorigenesis. In addition, PROX1 was also significantly increased in a FP-RMS model induced by forced PAX3-FOXO1 expression in mesenchymal stem cells (Ren et al., 2008). In both of the aforementioned RMS models of non-myogenic origin, GO analysis of the upregulated genes showed overrepresentation of muscle contraction-related genes, while the cell adhesion was overrepresented among the downregulated genes. Importantly, this is exactly the opposite to what we observed in PROX1 silenced RD cells, demonstrating that PROX1 silencing in RMS cells transforms the transcriptome to resemble that of benign mesenchymal stem cells. Our current findings in combination with the analyses of the publicly available data from previous studies indicate that PROX1 is needed for the muscle specification program in RMS, irrespective of the origin of the RMS. Moreover, our work strongly suggests that PROX1 silencing in RMS not only reverts the muscle phenotype and cell adhesion properties, but allows the RMS cells to regain characteristics of their cell of origin. This could provide insight in which cell lineages contribute to RMS tumorigenesis and the underlying mechanisms.

In previous studies, PROX1 expression has been associated with tumor progression and prognosis in other cancer types (Elsir et al., 2012). For instance, we have shown that PROX1 is important in the transition of benign colon adenoma to carcinoma, and in the maintenance of cancer stem cell features in intestinal adenomas and colorectal cancer (Petrova et al., 2008; Wiener et al., 2014). PROX1 is also essential for the clonogenic growth in colorectal cancer cells (Ragusa et al., 2014) and high PROX1 expression is correlated with a poor prognosis of esophageal cancer (Yokobori et al., 2015) and rectal neuroendocrine tumor patients (Jernman et al., 2015). However, the role of PROX1 in RMS development, growth or self-renewal has not been previously reported.

Histone acetylation in an important regulator of transcription, and HDAC inhibitors have been shown to induce cell death and suppression of RMS cell growth *in vitro* (Kutko et al., 2003). Recently, a class I HDAC inhibitor entinostat was shown to inhibit RMS tumor growth in preclinical tumor models (Bharathy et al., 2019; Bharathy et al., 2018). Furthermore, it was shown that HDACs are essential for the expression of RMS core regulator transcription factors, and that these core transcription factors were downregulated by class I HDAC inhibitors (Gryder et al., 2019). Importantly, PROX1 was among these core factors. In colorectal cancer, PROX1 was shown to maintain cancer stem cells by recruiting the NuRD complex that suppressed the Notch pathway activity (Hogstrom et al., 2018). Akin to this observation, our work uncovers that PROX1 interacts with the NuRD and CoREST deacetylase complexes in RMS. The transcriptional repressors HDAC1 and HDAC2 are the core proteins in these complexes. We observed similar transcriptional effects of PROX1 and HDAC1 silencing in the RMS cells, indicating that PROX1 and HDACs function in a complex to regulate the expression of RMS oncogenes. Because HDAC inhibitors have demonstrated efficacy in preclinical RMS models, and some of them have entered early-phase clinical trials for other pediatric cancers (clinicaltrials.gov), this further emphasizes the potential of interference of the PROX1-HDACs interaction in the treatment of RMS. More specifically, inhibiting the CoREST complex could be an attractive target in the future (Kalin et al., 2018).

Because solid tumors use *de novo* lipid synthesis to maintain supply of energy for growth and for membrane synthesis, a high rate of lipogenesis is induced in cancer cells (Mounier et al., 2014). Lipid oxidation has been shown to provide high energy required for tumor cell migration and metastasis in ARMS (Liu et al., 2012). PROX1 in lymphatic endothelial cells increases fatty acid oxidation during cell proliferation, and by interacting with p300 provides epigenetic regulation of lymphangiogenic gene expression (Wong et al., 2017). In hepatocytes, HDAC3–PROX1 interaction was found to be a major regulator for genes related to lipid homeostasis, and specific ablation of either component increased triglyceride content in the liver (Armour et al., 2017). In agreement with these results, PROX1 interacted with HDAC3 in RD cells, and PROX1 silencing led to lower fatty acid uptake and oxidation, accompanied by triglyceride accumulation, indicating that PROX1 also regulates tumor lipid metabolism. Moreover, transcriptomic analysis revealed that pathways related to cholesterol biosynthesis, isopropanoid biosynthesis, steroid biosynthesis and fatty acid metabolism were significantly repressed in the PROX1 silenced RD cells. PROX1 silencing also led to coordinately repressed expression of genes involved in oxidative phosphorylation, and forced the cells to utilize glycolysis instead of oxidative metabolism. Interestingly, in colorectal cancer, PROX1 promoted metabolic adaptation for maintained cell growth by enhancing glycolytic or oxidative metabolism in a context-depended manner (Ragusa et al., 2014).

In conclusion, we identified PROX1 as an important regulator of the muscle specification program, tumor growth and stemness as well as tumor cell metabolism in RMS. Mechanistically, PROX1 acts in NuRD and CoREST complexes with HDACs and regulates the expression of genes previously shown to influence RMS growth and cancer metabolism. Transcription factors have traditionally been considered as undruggable, but transcriptional regulation is emerging as a potential therapeutic target (Bhagwat & Vakoc, 2015). Recently, it was demonstrated that targeting the transcription factor-associated epigenetic machinery could be an effective way to inhibit RMS cell growth (Gryder et al., 2019). Our work increases understanding on the mechanisms regulating the myogenic phenotype and development of RMS, and suggests that targeting the PROX1-HDAC interaction could provide novel treatment options for RMS.

## Acknowledgements

We would like to thank Dr. Seppo Kaijalainen for cloning the BioID vectors and Dr. Jenny Högström-Stakem for discussions and comments during the project. Kirsi Mattinen, Maxime Laird, Manon Gruchet, Ilse Paetau, Tanja Laakkonen and Tapio Tainola are acknowledged for their excellent technical help. We also thank the Laboratory Animal Center at the University of Helsinki for expert animal care, the Biomedicum Imaging Unit for microscope support, the Biomedicum Functional Genomics Unit for the RNAseq experiments and the FIMM Technology Centre High Throughput Biomedicine for the drug sensitivity and resistance testing. Our first findings and analyses of this study were made in the Translational Cancer Biology Program, University of Helsinki and Wihuri Research Institute. The work was funded by the Cancer Foundation Finland, the Academy of Finland (grants 297245, 320185, 292816, 273817 and 307366), the Sigrid Jusélius Foundation, Children’s Cancer Foundation Väre, the Doctoral School of Biomedicine, iCAN Digital Precision Cancer Medicine Flagship, Helsinki Institute of Life Sciences (HiLIFE) and Biocenter Finland.

## Author contributions

N.Y.G. K.A. and R.K. conceptualized the study, N.Y.G. and R.K. designed and N.Y.G. performed the experiments. P.K. and T.P. assisted with experimental design and execution of the experiments. M.K. and M.V. performed the mass spectrometry analyses. E.G. and K.L. provided FGFR4 reagents and assisted with FGFR4 analyses. C.H., T.B. and M.S. provided the patient tumor samples and their IHC staining and participated in the analyses. R.K. supervised the project. N.Y.G., P.K., K.A. and R.K. wrote and all authors commented on the manuscript.

## Materials and methods

### Human RMS cell lines

Human RD cells were obtained from ATCC (CCL-136). KLHEL1 cells were derived from a biopsy of a 27-year-old male, diagnosed with high-grade alveolar RMS, FISH positive for FKHR (FOXO1) 13q14 gene fusion and positive for vimentin, MYF4, myosin and desmin. The ethical committee of the Helsinki University Hospital approved the study and written consent was obtained from the patient. RH30, RMS-YM and RH36 cell lines were kindly provided by Dr. Monika Ehnman (Karolinska Institute, Sweden). All cell lines were authenticated by STR and SNP profiling using ForenSeq DNA signature kit (Verogen) prior to use and tested regularly for possible mycoplasma contamination.

### Cell culture

The cells were cultured in Dulbecco’s Modified Eagle Medium (DMEM high glucose) supplemented with 10% fetal bovine serum (FBS), 2mM L-glutamine, penicillin (100U/ml) and streptomycin (100U/ml) and maintained at 37°C in a humidified 5% CO_2_ incubator. The lentiviral PROX1 silencing constructs (shPROX1 (Wiener et al., 2014) TRCN0000016252), FGFR4 silencing constructs (TRCN0000000628 and TRCN0000010531), shHDAC1 silencing constructs (TRCN0000004817, TRCN0000004818) and the scramble control (SH002) were from the TRC library. The cells were transduced with 293FT supernatant containing lentivirus for two to three days. For proliferation analysis, 1 × 10^5^ cells were seeded on coverslips and cultured overnight. Next day the cells were incubated four hours with 10nM EdU and the rest of the procedure followed Click-iT^®^ EdU imaging Kit Protocol (Thermo Fisher Scientific). For the clonogenic assay, 2000 RD cells and 500 KLHEL1 cells were plated in 6-well dish and cultured for 13 days before analysis. Colony number was measured with GelCount™ (Oxford Optronix). For spheroid analysis, 2000 RD cells and 500 KLHEL1cells were embedded in 15 µl of Matrigel (Corning) and analyzed at day nine. Spheroids were imaged with EVOS FL inverted epifluorescence microscope (Thermo Fisher Scientific) using 4X or 10X LD Ph Air objectives.

### Bioinformatics analysis of human RMS datasets

Publicly available data sets from previous studies on RMS tumors deposited in Gene Expression Omnibus (https://www.ncbi.nlm.nih.gov/geo/), cBioportal (https://www.cbioportal.org/) and Oncomine (https://www.oncomine.org/resource/login.html) were analyzed for PROX1 expression. The following data sets were used (GEO accession number): GSE108022, GSE22520, GSE2553.

### BirA-mediated proximity labeling

About 10 million RD cells per sample were transduced with or FUW-BirAv-hPROX1 or FUM-BirAv-NLS-Cherry. Next, cells were incubated with 50 mmol/L biotin (Sigma Aldrich #B4501) for 24h, washed twice with cold PBS, collected with PBS plus protease inhibitors, and stored in −80 ° C. The cells were lysed in BioID – Lysis buffer (HENN – buffer with 0.5 % IGEPAL, 0.1% SDS, 1mM PMSF, 1.5 mM Na3VO4, 1mM DTT, Sigma Protease inhibitor cocktail) with 1 µL Benzonase Nuclease by incubating on ice for 10min, followed by three cycles of sonication and centrifugation twice at 16,000. Insoluble materials were discarded and clear lysates were added to Strep-Tactin sepharose (400 μL 50% slurry; IBA Lifesciences #2-1201-010) loaded columns (Bio-Rad Laboratories #732-6008) and washed with ice cold HENN – buffer. Proteins were eluted with 0.5mM biotin - HENN – buffer. The remaining procedure and liquid chromatography–tandem mass spectrometry analysis were conducted as described previously (X. Liu et al., 2018)

### BioID data analysis

The data from the PROX1 BioID MS analyses (2 biological and 2 technical replicates), were filtered by retaining only preys that were identified in all four replicates. CRAPome contaminant database was used to screen out the common contaminants seen in more than 25% of the control analyses, but preys with at least double the PSM value compared to the corresponding CRAPome average PSM value, were kept in the filtered data set.

### Mouse tumor models

For xenograft experiments, 8-12-week old NOD *scid* gamma (NSG, NOD.Cg-Prkdc scid Il2rg tm1Wjl/SzJ, 005557) female mice from Jackson laboratory were used. 2 × 10^6^ shProx1 or shSCR RD cells were embedded into 100 µl Gibco Geltrex™ and injected subcutaneously into the right and left scapular area of isoflurane anesthetized NSG mice. The mice were checked regularly for palpable tumors. Tumor growth was monitored weekly by using calipers. Measurements of the height (H), width (W), and depth (D) were taken and converted into relative tumor volume. At pre-defined time points, the mice were euthanized and tumors harvested. Volume and mass of the excised tumors were measured, and the tumor was subjected for histological and gene expression analysis. All animal experiments were approved by the National Animal Experiment Board in Finland (ESAVI/6306/04.10.07/2016).

### Histology and immunohistochemistry

Harvested tumors were fixed in 4% paraformaldehyde (PFA), processed and embedded into paraffin. 5 µm sections were deparaffinized and stained with either hematoxylin and eosin or with specific antibodies for PROX1 (R&D Systems, AF2727) and GFP (Abcam, Ab13970). After deparaffinization, sections were subjected to heat-induced epitope retrieval (High pH Retrieval solution, DAKO). For signal detection, Alexa Fluor 488 and 594 -conjugated secondary antibodies (Molecular Probes, 1:500) were used. PROX1 staining of the patient RMS samples was performed by using the BenchMark XT automated system on 5 µm thick, antigen-retrieved sections. Cultured cells were fixed with 4% PFA and permeabilized and blocked for nonspecific antibody binding with 0.3% Triton X-100 and 3% BSA. Overnight incubation with primary antibodies based on manufactures dilution recommendations was followed by staining with Alexa Fluor-conjugated secondary antibodies (1:500) and DAPI for nuclear staining. Samples were imaged using Leica DM LB light microscope, Zeiss Axioplan fluorescent microscope and Zeiss LSM 780 confocal microscope. Images were processed and analyzed with ImageJ (NIH).

### RNA extraction and quantitative real time PCR

Total RNA from the lysed cells was extracted with Nucleospin RNA II Kit (Macherey-Nagel). The tumors were first lysed with Trisure reagent (Bioline) and RNA was further purified with the Nucleospin RNA II Kit (Macherey-Nagel). cDNA synthesis was performed with the High-Capacity cDNA Reverse Transcription Kit (Thermo Fisher Scientific) using 1 mg of RNA and random hexamer Primers. qRT-PCR was performed with Fast Start Universal SYBR Green Master Mix (Applied Biosystems). Relative gene expression levels were calculated using the formula 2 ^−CtΔΔ^. All primer sequences are shown in Supplementary Table 3.

### Protein extraction, Immunoprecipitation and western blot analysis

Total protein from human RMS cells was obtained by lysis in RIPA buffer (50mM Tris-HCl at pH7.6, 150mM NaCl, 1% NP-40, 0.5% DOC, 0.1% SDS, with protease and phosphatase inhibitor coctail (Pierce)). The concentration of the lysates was measured with the BCA protein assay kit (Thermo Fisher Scientific) following the manufacturer’s instructions. The samples were loaded into 4%–20% Mini-Protean TGX gels (Bio-Rad Laboratories), transferred onto nitrocellulose membranes and incubated overnight at +4 with primary antibodies against PROX1 (goat-anti-human, AF2727, R&D, 1:1000) and vinculin (mouse anti-vinculin, V9131, Sigma, 1:1000). HRP conjugated secondary antibodies (Dako, 1:5000) were used. The signal was visualized by the SuperSignal West Pico or Femto Maximum Sensitivity Substrate (Thermo Fisher Scientific) and detected with LiCor Odyssey®. For co-immunopreciptation (co-IP), the RD cells were pretreated with 1.5 mM amine reactive cross-linker DSP (Thermo Fisher Scientific) and lysed with IP lysis buffer containing protease inhibitors (Pierce). The IP was performed using Dynabeads® according to the manufacturer’s protocol. Antibodies against PROX1 (11067-2-AP, Cell Signaling), HDAC1 (5356, Cell Signaling), LSD1 (ab17721, Cell Signaling) and MTA1 (5646, Cell Signaling) were used.

### RNA sequencing and enrichment analysis

1 μg of total RNA was treated with Ribo-Zero Complete Gold Kit to remove the ribosomal RNA and purified with Qiagen RNeasy MinElute Cleanup Kit. The absence of rRNA and the quantity was determined by Bioanalyzer. NEBNext Ultra Directional RNA Library Prep Kit for Illumina was used to generate cDNA libraries for next generation sequencing. The library quality was assessed by Bioanalyzer (Agilent DNA High Sensitivity chip) and quantity by Qubit (Invitrogen). The analysis from the generated FASTQ data was performed using Chipster software (www.chipster.csc.fi) (Kallio et al., 2011). In brief, the quality of FASTQ files were assessed with FASTQC, and used as input for alignment to reference genome using HISAT2 (Kim, Langmead, & Salzberg, 2015). Aligned reads per gene were counted with HTseq (Anders, Pyl, & Huber, 2015) and DEseq2 (Love, Huber, & Anders, 2014) was used to anayze differentially expressed genes. Fragments Per Kilobase of transcript per Million mapped reads (FPKM) values were used and statistical significance of adjusted P-value (FDR) < 0.05 was set as cut off, and the final values were log2 transformed. The differentially expressed genes with fold change of log2 ≤ −0.25 and log2 ≥ 0.25 were transferred to Gene Set Enrichment Analysis software (GSEA, http://www.broadinstitute.org/gsea) (Mootha et al., 2003; Subramanian et al., 2005) to analyze enrichment for curated gene sets (ftp.broadinstitute.org://pub/gsea/gene_sets/c2.all.v7.0.symbols.gmt). The analysis was performed with default parameters employing GseaPreranked tool with 1000 permutations and gene sets of maximum size of 500 and minimum 15. Similarly, gene ontology (GO) and pathway enrichment analyses were performed by using annotation Database, Visualization and Integrated Discovery (DAVID software; http://david-d.ncifcrf.gov) (Huang da, Sherman, & Lempicki, 2009). Benjamini-Hochberg method was implemented to adjust P-value.

### Cell viability and cytotoxicity assays

MTT assay was used to test the cytotoxicity of Vorinostat. ShSCR and shPROX1 RD cells were seeded on 96-well plates at concentration 5000 cells/well and incubated for 24 hours. The following day, cells were either treated with the drugs or vehicle (DMSO). After 48 hours of treatment, 10 µl of 12mM MTT stock solution (Thiazolyl Blue Tetrazolium Bromide, Sigma) was added to all wells and left for 2-4 hours in 37°C until formation of formazan crystals. Then, medium was removed, cells were washed two times with PBS and formazan was solubilized by adding 100 µl of DMSO. Absorbance was measured at 540 nm using the FLUOstar® Omega (BMG Labtech).

### Statistical analyses

All statistics analyses except for RNA sequencing were done using GraphPad Prism (V.7.0). Unpaired two-tailed *t*-test was performed for experiments with two groups. Statistical significance levels were defined as \**P* < 0.05; \*\**P* < 0.01; \*\*\**P* < 0.001. One-way ANOVA with Tukey’s multiple comparisons test was used for more than two groups.

## Supplemental Information titles

**Supplement Table 1.** Downregulated and Upregulated genes in PROX1 silenced RD cells

**Supplement Table 2.** Proteins identified to interact with PROX1 by PROX1–BioID fusion protein expression and mass spectrometry analysis of the biotinylated proteins.

**Supplement Table 3.** Primers for qPCR

---

**Supplemental Figure 1.**
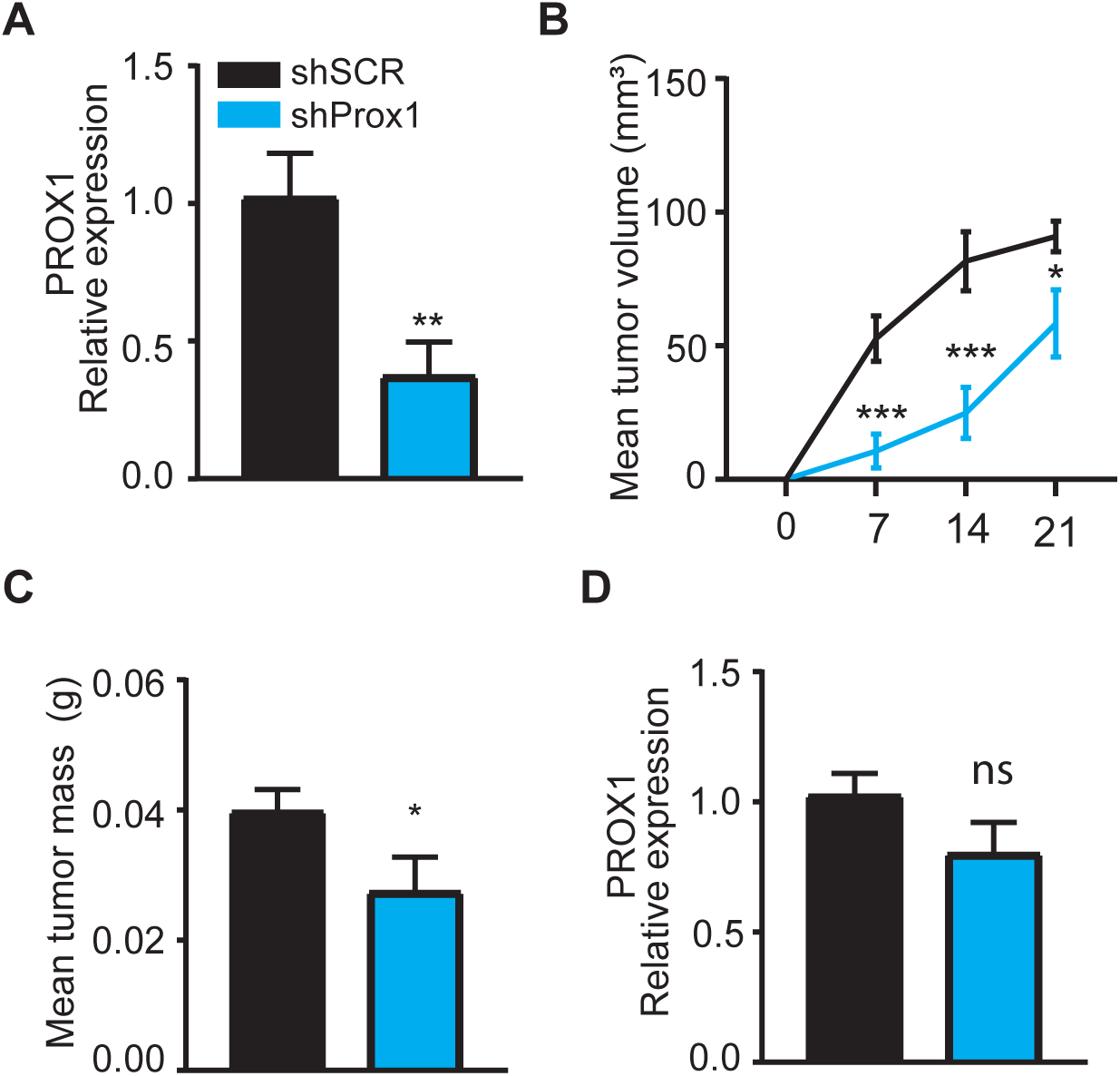
PROX1 regulates the growth of RMS xenograft. (**A**) RT-qPCR analysis of the PROX1 mRNA expression in shPROX1 (construct 2) RD cells before tumor implantation. (**B**) Quantification of tumor growth based on tumor volume (mm^3^) for shSCR and shPROX1 (2) RD cells injected into NSG mice left and right flank region, respectively (n = 9 per group). (**C**) Comparison of average tumor mass at the end of the experiment (day 21). (**D**) RT-qPCR analysis of PROX1 expression in harvested tumors. *P <0.02 and **P<0.01; n.s., not significant. Data is presented as mean ± SEM.

**Supplemental Figure 2.**
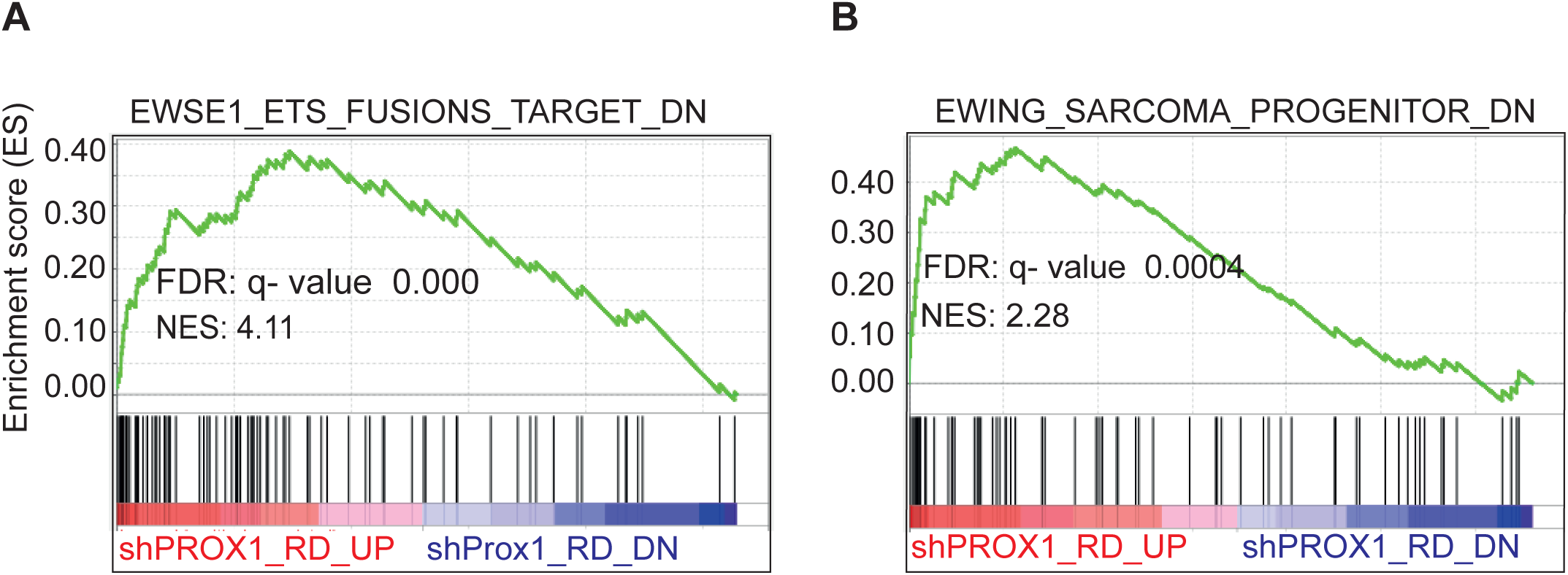
PROX1 regulate malignant transcriptomic phenotype of RMS and EFT. (**A and B**) Gene Set Enrichment Analysis plots for gene sets overlapping with PROX1 driven differentially expressed genes in RD cells. (**A**) Enrichment of EWS/FLI1 and EWS/ERG fusion gene targets in mesenchymal progenitor cells. (**B**) Enrichment of EWS-FLI1 fusion gene targets in mesenchymal stem cells (Ewing sarcoma progenitors). NES, normalized enrichment score; FDR, false discovery rate. Each black bar represents a single gene within a gene set.

**Supplemental Figure 3.**
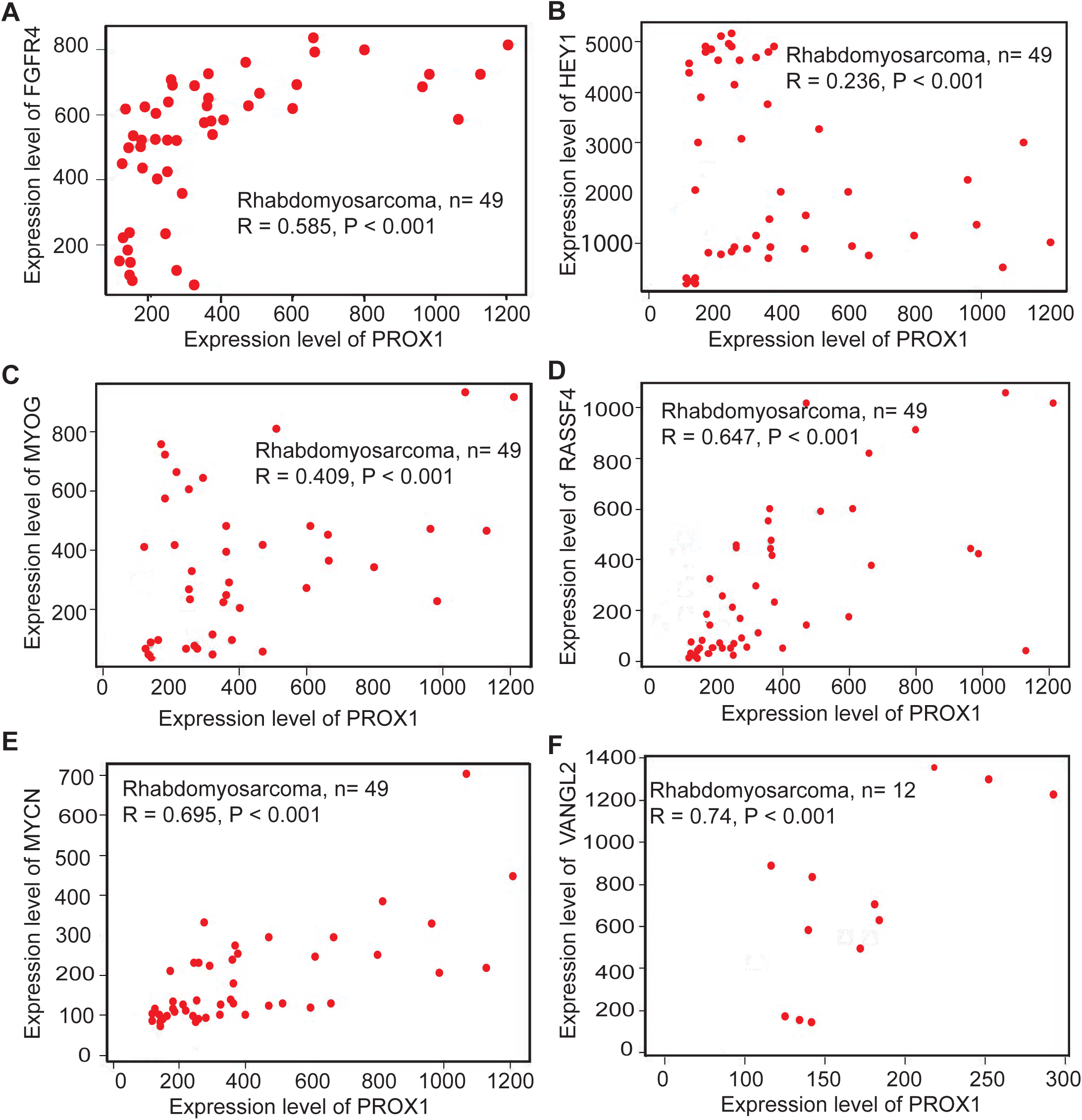
PROX1 mRNA expression correlates with several RMS oncogenes in RMS tumors. (**A-F**) Correlations between PROX1 and known RMS pro-oncogenic genes in RMS tumors, analyzed using Medisapiens database.

**Supplemental Figure 4.**
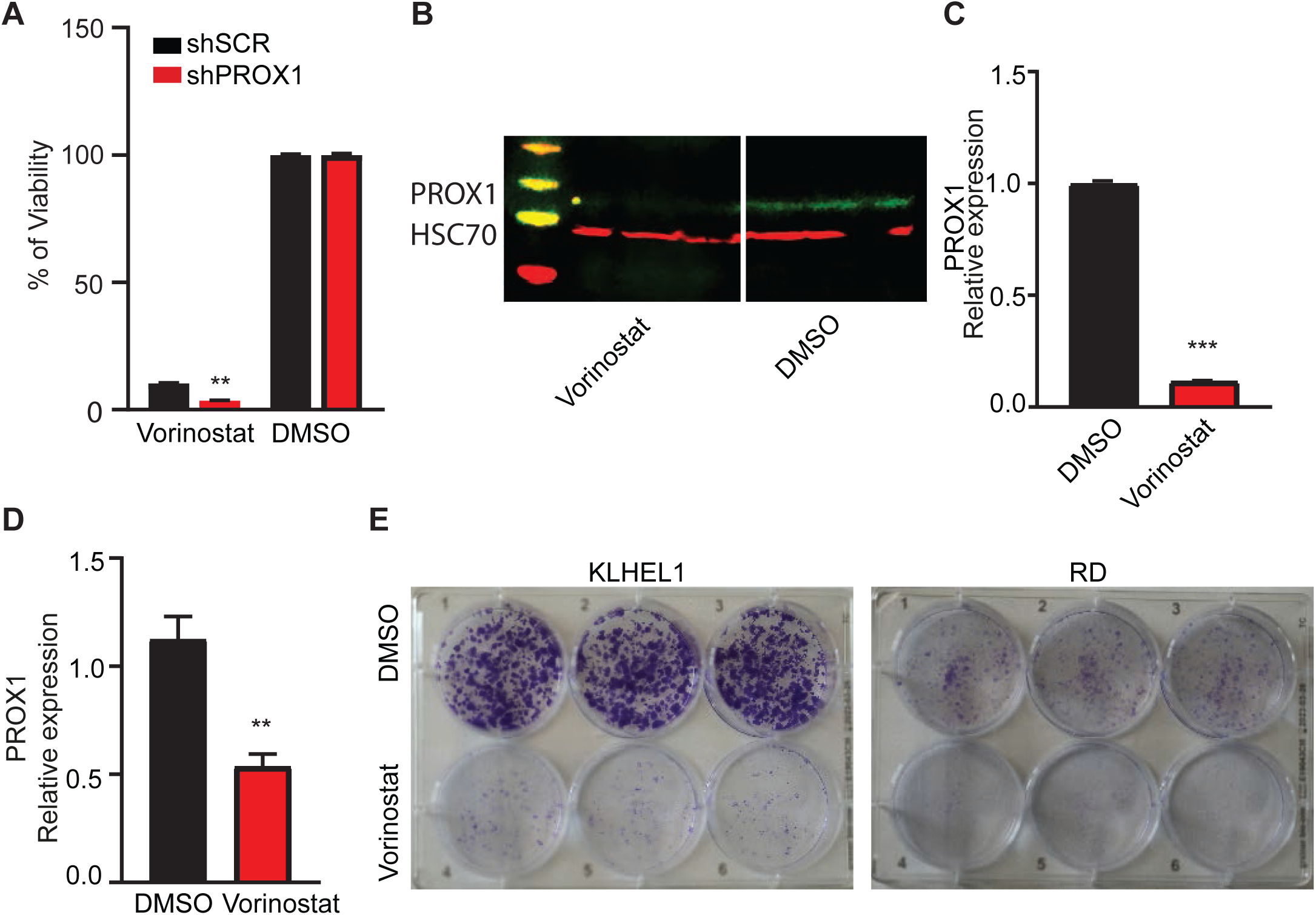
PROX1 silencing enhances the effect of HDAC inhibitor Vorinostat on RMS cells. (**A**) Cell viability assay of shSCR and shPROX1 RD cells treated with Vorinostat and DMSO as a control. (**B**) Western blot analysis of PROX1 expression in Vorinostat treated RD cells. HSC70 was used as loading control. (**C**) Quantification of western blot. (**D**) RT-qPCR analysis of the PROX1 mRNA expression in DMSO and Vorinostat treated RD cells. (**E**) Colony formation assay in KLHEL1 and RD cells treated with Vorinostat. 1000 cells for KLHEL1 and RD were plated per well and get fixed and stained with crystal violet 9 days after seeding. **P <0.01 and ***P<0.001. Data is presented as mean ± SEM.

**Table S3.**
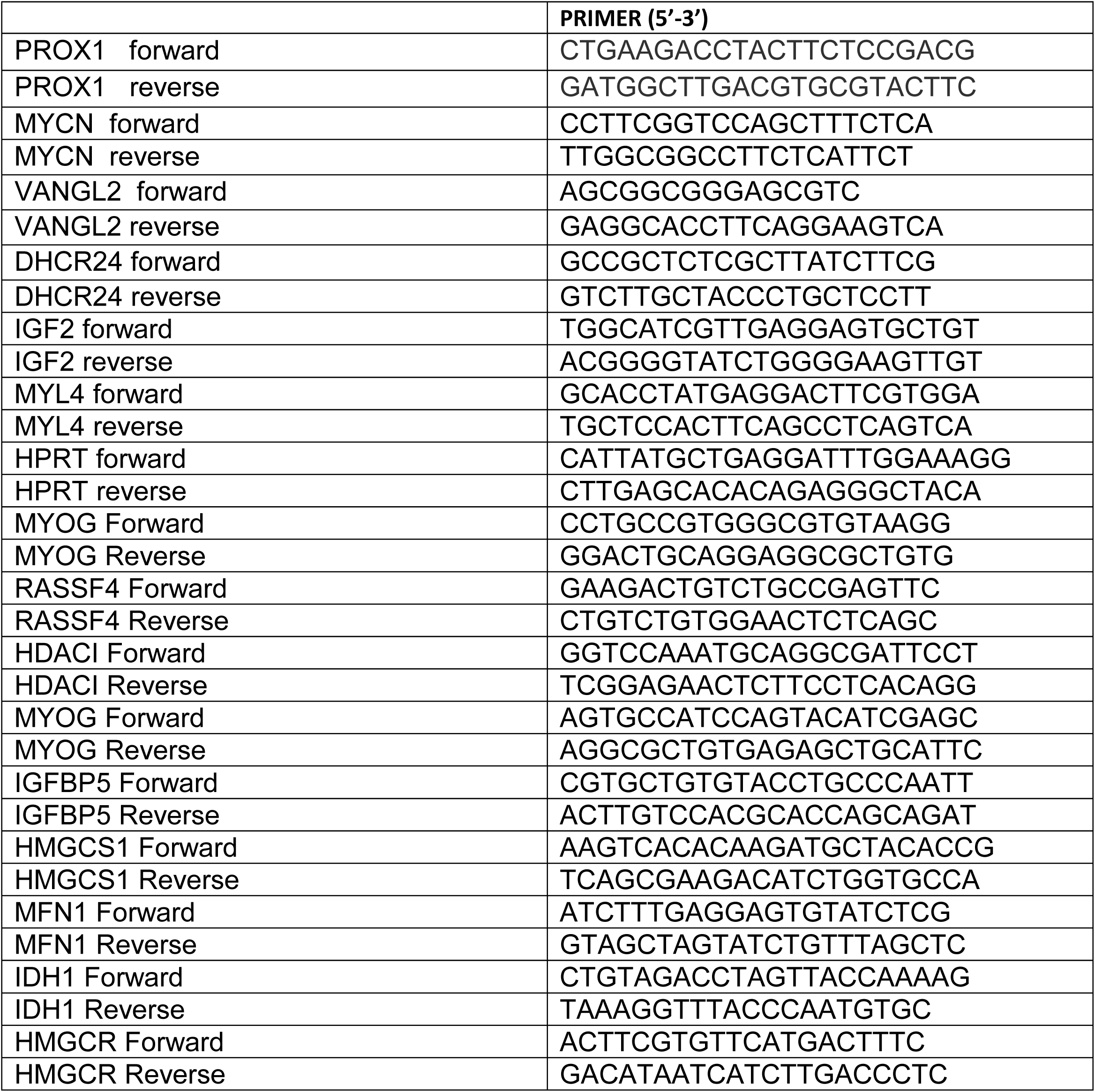
Human qPCR SYBR Green primers.

